# 2,4-Di-tert-butylphenol Induces Adipogenesis in Human Mesenchymal Stem Cells by Activating Retinoid X Receptors

**DOI:** 10.1101/2022.10.08.511439

**Authors:** Xiao-Min Ren, Richard C. Chang, Yikai Huang, Angélica Amorim Amato, Coralie Carivenc, Marina Grimaldi, Angela Y. Kuo, Patrick Balaguer, William Bourguet, Bruce Blumberg

**Affiliations:** Department of Developmental and Cell Biology, University of California, Irvine, CA, USA; Department of Pharmaceutical Sciences, University of California, Irvine, CA, USA; Department of Biomedical Engineering, University of California, Irvine, CA, USA; Centre de Biologie Structurale, Université Montpellier, CNRS, Inserm, Montpellier, France; Institut de Recherche en Cancérologie de Montpellier (IRCM), Inserm U1194, Université Montpellier, Institut régional du Cancer de Montpellier (ICM), Montpellier, France; Faculty of Environmental Science and Engineering, Kunming University of Science and Technology, Kunming 650500, China

**Keywords:** Endocrine disruptor, EDC, RXR, PPAR, MSC, obesogen, adipogenesis

## Abstract

2,4-di-tert-butylphenol (2,4-DTBP) is an important commercial antioxidant and a toxic natural secondary metabolite that has been detected in humans. However, there is scant information regarding its toxicological effects. Here we asked whether 2,4-DTBP is a potential obesogen. Using a human mesenchymal stem cell (MSC) adipogenesis assay, we found that exposure to 2,4-DTBP led to increased lipid accumulation and expression of adipogenic marker genes. Antagonist assays revealed that 2,4-DTBP increased lipid accumulation by activating the peroxisome proliferator-activated receptor γ (PPARγ)-retinoid X receptor (RXR) heterodimer. 2,4-DTBP likely activated the PPARγ/RXRα heterodimer by activating RXRα but not directly binding to PPARγ. We confirmed that 2,4-DTBP directly bound to RXRα by solving the crystal structure of this complex, then predicted and demonstrated that related compounds could also activate RXRα. Our study demonstrated that 2,4-DTBP and related chemicals could act as obesogens and endocrine disruptors via RXR. These data showed that 2,4-DTBP belongs to a family of compounds whose endocrine-disrupting and obesogenic effects can be strongly modulated by their chemical composition and that structure-activity studies such as the present one could help guide the rational development of safer antioxidants.

**SYNOPSIS:** Little research exists on the effects of commercially valuable antioxidants on biological systems. This study reports that di- and tri-tert-butylphenols can act as endocrine disruptors and potential obesogens by activating nuclear hormone receptors.

## INTRODUCTION

Obesity has become a serious health problem worldwide in infants, children, adolescents and adults ^1^. Approximately 13% of the world population is clinically obese (body mass index, BMI ≥ 30 kg/m^2^) and 39% of adults are overweight (BMI ≥25 kg/m^2^) ^2^. Obesity can increase the risk of many chronic diseases such as type 2 diabetes, heart disease, hypertension, stroke, and certain cancers ^3^ and increases risk of serious illness from COVID-19. According to the World Health Organization, at least 2.8 million deaths annually could be related to overweight or obesity worldwide ^4^. Obesity is an even greater problem in the U.S. where 42% of adults are currently obese ^5^.

Obesity is considered to be a result of the combined effects of both genetic and environmental factors ^6^. Traditional risk factors include lack of physical activity, excessive consumption of food, genetic susceptibility, and epigenetic predisposition ^7^. However, these factors cannot fully explain the dramatic increase in the obesity rate worldwide. An increasing number of epidemiological studies together with experimental evidence in animal models indicated that some environmental chemicals might be important contributors to obesity ^6^. Such “obesogens” include natural chemicals, pharmaceutical chemicals, or xenobiotic chemicals that could promote obesity ^8^. Known obesogens include organotins, bisphenols, agrochemicals, perfluoroalkyl substances, phthalates, and polybrominated diphenyl (reviwed in ^9^). Obesogens can contribute to obesity by increasing white adipocyte cell number or increasing lipid storage in existing adipocytes ^10^. These chemicals might also lead to obesity by affecting appetite and satiety, changing basal metabolic rate, altering energy balance, or influencing gut microbiota ^6^. Identifying which environmental pollutants are obesogens and revealing the underlying mechanisms could help to ameliorate the obesity pandemic and related diseases.

2,4-Di-tert-butylphenol (2,4-DTBP) is an important commercial antioxidant that is mainly used as an intermediate to produce other high molecular weight antioxidants and UV stabilizers for hydrocarbon-based products and plastics ^11^. 2,4-DTBP was detected in indoor dust ^12^, river sediment ^13^, wastewater, and sludge ^14^. 2,4-DTBP is also a natural toxic secondary metabolite produced by bacteria, fungi, plants, and animals ^15^. Human exposure can result from packaged food intake because tris(2,4-di-tert-butylphenol) phosphite in food packaging materials can break down to 2,4-DTBP ^16^, ingestion of dust ^12^ and intake from food containing 2,4-DTBP ^17^. A recent study analyzing synthetic phenolic antioxidants (SPAs) in humans detected 2,4-DTBP in urine samples from U.S. ^18^. 2,4-DTBP was detected in 100% of the investigated human urine samples and was the predominant congener, contributing 88.2% to total target concentrations of SPAs in the urine samples. The concentration of total 2,4-DTBP [7.3-130 ng/mL (0.03 μM-0.63 μM), geometric mean (GM): 25.8 ng/mL; 0.12 μM] was much higher than 2,6-di-tert-butyl-4-methylphenol (BHT) (0.85 ng/mL, 3.8 nM) and 3-tert-butyl-4-hydroxyanisole (BHA) (0.39 ng/mL, 2 nM), which were two other SPAs of concern. The relatively high levels of 2,4-DTBP found in human urine led us to investigate its possible adverse impacts.

In the present study, we assessed the capability of 2,4-DTBP to induce adipogenesis in human multipotent mesenchymal stromal stem cells (a.k.a. mesenchymal stem cells or MSCs) and explored its mechanism of action. We found that 2,4-DTBP, but not its structural analog 2,6-di-tert-butylphenol (2,6-DTBP), which is also used industrially as a UV stabilizer and antioxidant for hydrocarbon-based products ^11^, could induce adipogenesis and expression of white adipocyte marker genes in MSCs. We used receptor selective agonist and antagonist studies to link 2,4-DTBP exposure with activation of the peroxisome proliferator activated receptor gamma (PPARγ) and its heterodimeric partner, the retinoid X receptor (RXR) to promote adipogenesis in MSCs.

Reporter gene assays revealed that 2,4-DTBP but not 2,6-DTBP could activate RXRα, the RXRα subunit of the PPARγ/RXRα heterodimer and RXRα in the context of two other nuclear receptor complexes: the liver X receptor alpha (LXRα)/RXRα heterodimer and the thyroid hormone receptor β (TRβ)/RXRα heterodimer. We solved the crystal structure of 2,4-DTBP complexed with RXRα and identified the mechanistic basis underlying binding of 2,4-DTBP to RXRα and activation of RXRα by 2,4-DTBP. This analysis suggested that other related chemicals such as 2,4,6-tri-tert-butyl phenol and 1,3,5-tri-tert-butylbenzene would activate RXRα whereas 2,6-DTBP and 1,3-di-tert-butylbenzene could not. Reporter gene assays confirmed these predictions. Overall, our studies identified 2,4-DTBP as an RXRα activator and therefore a potential endocrine disrupting chemical (EDC) and obesogen.

## METHODS

### Chemicals and reagents

Dexamethasone, isobutylmethylxanthine, Nile red, Hoechst 33342, HX531, LG100268 (LG), 2,4-di-tert-butylphenol (2,4-DTBP), 2,6-di-tert-butylphenol (2,6-DTBP), 2,4,6-tri-tert-butylphenol (2,4,6-TTBP), 1,3-di-tert-butylbenzene (1,3-DTBB), 1,3,5-tri-tert-butylbenzene (1,3,5-TTBB), LG100268, GW3065, T3 and TTNPB were purchased from Sigma-Aldrich. CD3254 was purchased from Tocris Bioscience. Rosiglitazone (ROSI) was purchased from Cayman Chemical Company. T0070907 was from Enzo Life Sciences. Dimethylsulfoxide (DMSO) was purchased from Thermo Fisher Scientific.

### Human MSCs adipogenesis assay

Human bone-marrow MSCs (MSCs) were purchased from PromoCell at passage 2 (Lot: 429Z001-Female). Human MSCs were cultured in alpha Modification of Eagle’s minimal essential medium (αMEM) (Thermo Fisher Scientific) supplemented with 15% FBS (Gemini Bio-Products), 2% penicillin, 5,000 IU/mL-streptomycin 5,000 μg/mL (Corning), 10 μM HEPES (Fisher Chemical). The cells were split when the confluency reached 90% and used at passage 6 for adipogenesis assays. 40,000 cells per well were seeded in a 24-well plate or 80,000 cells per well were seeded in a 12-well plate. When cells reached 100% confluency, media were replaced with differentiation media [αMEM, 15% FBS supplemented with adipogenic induction cocktail (MDI: 500 μM isobutylmethylxanthine, 1 μM dexamethasone, 5 μg/mL human recombinant insulin] and ligands. Specific ligands (dissolved in DMSO) were administered twice weekly for 2 weeks. Antagonist assays were performed similarly, except that T0070907 was added twice daily (to 10 μM). DMSO concentrations in the medium were identical between vehicle controls and test chemicals and never exceeded 0.1%. At the end of each assay, cells in the 24-well plates were fixed in 3.7% formaldehyde in PBS for 30 min at 25°C for lipid staining; cells grown in 12-well plates were homogenized and total RNA extracted for gene-expression analysis.

### Gene expression analysis

Total RNA was isolated from cells using Trizol reagent as recommended by the manufacturer (Invitrogen Life Technologies). Reverse transcription and QPCR were performed using Transcriptor reverse transcriptase and Sybr Green Master Mix (Roche Diagnostics Corp., Indianapolis, IN). The following genes were examined: FABP4 (fatty acid binding protein 4), FSP27 (fat-specific protein of 27 kDa), PPARγ2 and LPL (lipoprotein lipase). The details of the experiment and sequences of primers used for QPCR were as described in our previous studies ^19, 20^.

### Transient transfection and luciferase reporter gene assays

The CMX-GAL4 effector plasmids using pCMX-GAL4 DNA-binding domain fusion constructs to nuclear receptor ligand-binding domains for human PPARγ (pCMX-GAL4-PPARγ), human RXRα (pCMX-GAL4-RXRα), human LXRα (pCMX-GAL4-LXRα), and human TRβ (pCMX-GAL4-TRβ) were previously described ^21-23^. Transient transfections were performed in COS-7 cells (ATCC CRL-1,651TM) as described ^20^. Briefly, COS-7 cells were seeded at 15,000 cells/well in 96-well tissue culture plates in Dulbecco’s Modified Eagle Medium (DMEM; HyClone), supplemented with 10% calf bovine serum (Atlanta Biologicals). 24 hours later, cells were transfected in Opti-MEM reduced-serum medium (Invitrogen Life Technologies, Grand Island, NY) when they reached ∼ 90% confluency (∼24 h after seeding). Cells were transfected 1 μg of CMX-GAL4 effector plasmids (0.3 μg for pCMX-GAL4-PPARγ plasmid) with or without 1 μg pCMX-ligand-binding domain of RXRα (L-RXRα) (0.3 μg pCMX-L-RXRα for pCMX-GAL4-PPARγ plasmid) was co-transfected with 5 μg tk-(MH100)4-luciferase reporter and 5 μg CMX-β-galactosidase transfection control plasmids in Opti-MEM using Lipofectamine 2000 reagent (Invitrogen Life Technologies, Grand Island, NY), following the manufacturer’s recommended protocol. After 24 h incubation, the medium was replaced with phenol red-free Dulbecco’s modified Eagle medium (DMEM) (Invitrogen Life Technologies) /10% resin charcoal-stripped fetal bovine serum (FBS) plus ligands at concentrations indicated in the figure legends for an additional 24 h before luciferase and β-galactosidase assays. 2,4-DTBP and 2,6-DTBP were tested from 1.2 μM to 40 μM with 2-fold between each other, with 40 μM producing no noticeable cytotoxicity as judged by reduced β-galactosidase activity (data not shown). The ligands were dissolved in DMSO. The DMSO concentrations in the medium were identical between vehicle controls and test chemicals and never exceeded 0.1%. 24 h after adding the ligands to the media, cells were lysed in 165 μL of lysis buffer and allowed to shake for 30 min at 25°C. For luciferase assay, 35 μL of the lysate from each well was transferred to a well in a clean, nontreated, white, flat-bottom, polystyrene, 96-well plate (Costar). Additionally, 100 μL of luciferase solution made fresh was added to each well with the cell lysate, and that plate was shaken for 30 s at RT. Plates were placed in Dynatech ML3000 luminometer, and data were acquired with luminometer ML3000 (version 3.07) software. For β-galactosidase assay, 35 μL of the cell lysate was transferred to a clear, flat-bottom, 96-well plate (Thermo Fisher Scientific). In addition, 100 μL of β-galactosidase solution was added to each well. Plates were shaken for 30 s and incubated at 25° C for 15 min. Absorbance was measured with a wavelength of 405 nm on a Versamax microplate reader (Molecular Devices) and SOFTmaxPRO 4.0 software. Each luciferase read was normalized with the corresponding β-galactosidase read coming from the same transfection well and multiplied by the number of minutes the β-galactosidase plate was incubated (15 min). Data are reported as relative light units (Luc/Gal). EC_50_ and maximum activations were calculated using GraphPad Prism 9.0 (GraphPad Software, Inc.). The transfections were performed in triplicate and reproduced in multiple experiments.

### Statistical analysis

The data are expressed as the mean value ± standard error of the mean (n=3). Unpaired t-test was used to determine the significance of effects elicited by the chemicals relative to the control group in human MSCs adipogenesis assays and to determine the significance of differences comparing +T0070907 versus –T0070907 or +HX531 versus –HX531 with each other. In the transfection assays, the significance of the experimental data was analyzed using one-way analysis of variance (ANOVA), followed by a least significant difference multiple comparisons test. A *p* value of less than 0.05 was considered to be statistically significant. Statistical analysis used GraphPad Prism 9.0 (GraphPad Software Inc., San Diego, CA).

### Retinoic acid receptor β (RARβ) reporter cell line

HELN RARβ cell line was already described ^24^. Briefly, we firstly generated the HELN cell line by transfecting HeLa cells with the p-ERE-βGlob-Luc-SVNeo plasmid. Secondly, we transfected HELN cells with pRARβ(ERαDBD)-puro plasmid. pERE-βGlob-Luc-SVNeo contains a luciferase gene driven by an estrogen receptor binding site in front of the β-globin promoter and a neomycin phosphotransferase gene under the control of the SV40 promoter. In pRARβ(ERαDBD), the encoded chimeric RARβ protein contains the DNA binding domain of the estrogen receptor α. Puromycin N-acetyl transferase selection marker expression confers resistance to puromycin.

To measure the activity of the chemicals, HELN RARβ cells were seeded at a density of 40,000 cells per well in 96-well white opaque tissue culture plates (Dutscher, Brumath, France). Compounds were added 8 h later alone in presence of the pan RAR-agonist TTNPB 100 nM or the pan RXR-agonist CD3254 100 nM and cells were incubated for 16 h. At the end of incubation, the culture medium was replaced by a medium containing 0.3mM luciferin. Luciferase activity was measured for 2 seconds in intact living cells using a microBeta Wallac luminometer (Perkin Elmer, Villebon sur Yvette, France). Tests were performed in quadruplicate in at least three independent experiments and data were expressed as mean ± s.e.m. Results are expressed in percentage of the maximal activity obtained in presence of TTNP 100 nM. Dose-response curves were fitted using the sigmoid dose–response function of a graphics and statistics software package (Graph-Pad Prism, version 4, 2003, Graph-Pad Software Inc., San Diego, CA, USA).

### Crystallographic analysis

The histidine-tagged LBD of human RXRα (residues 223-462 in a pET15b vector) was expressed in *Escherichia coli* BL21(DE3). Cells were grown in ZYM-5052 autoinduction medium supplemented with 100 µg/mL ampicillin. After 24 h incubation at 25°C, cell cultures were harvested by centrifugation at 8,000 × *g* for 20 min. The cell pellet from 2 liters of culture was resuspended in 250 mL buffer A (20 mM Tris-HCl pH 8.0, 500 mM NaCl, 5 mM imidazole) supplemented with 100 µg/mL of lysozyme and a protease inhibitor cocktail (Complete, mini, EDTA-free tablet, Roche Applied Science) then incubated with shaking for 30 min at 4°C. The suspension was lysed by sonication and centrifuged at 35,000 ×*g* for 30 min at 4°C. The supernatant was loaded onto a 5 mL Ni^2+^-affinity column (HiTrap chelating column, GE healthcare), preequilibrated with buffer A, using the Äkta pure system (GE Healthcare). The column was washed with 10 volumes of buffer A. Protein was eluted in 0-500 mM imidazole gradient using 25 column volumes of the buffer. The fractions containing RXRα LBD were pooled and incubated overnight at 4°C with thrombin to remove the histidine-tag. The protein was further purified using a Superdex 75 26/60 gel filtration column (GE healthcare) preequilibrated with buffer C (20 mM Tris-HCl pH 8.0, 150 mM NaCl, 5% Glycerol, 5 mM DTT).

Fractions containing the purified receptor were pooled, mixed with 1:3 molar ratio of 2,4-DTBP and 1:2 molar ratio of the TIF2 coactivator peptide and concentrated to 8 mg/mL. Crystals were obtained by vapor diffusion at 293° K by mixing 1 µL of protein with 2 µL of precipitant solution. The well buffer contained 20% PEG 3350 and 0.2 M sodium formate. A single crystal was mounted from the mother liquor onto a cryoloop (Hampton Research, Aliso Viejo, CA, USA), soaked in the reservoir solution supplemented with 20% glycerol and 40 nM of 2,4-DTBP and then frozen in liquid nitrogen. Diffraction data were collected on the ID30A-1 at the European Synchrotron Radiation Facility (Grenoble, France). Data were processed and scaled with XDS ^25^ and Scala ^26^. Crystals belong to space group P212121. The structure was refined using REFMAC5 ^27^ and COOT ^28^. Figures were generated with PyMol (https://pymol.org/2/).

## RESULTS

### 2,4-DTBP increased adipogenesis in human MSCs

MSCs have been widely used to study the obesogenic properties of environmental chemicals ^19, 20, 29, 30^. Here, we tested the potential obesogenic properties of 2,4-DTBP and 2,6-DTBP using a human MSC adipogenesis assay. The selective PPARγ agonist, ROSI was used as a positive control, DMSO vehicle (0.1% v/v) was the negative control. Treatment of MSCs with adipogenesis cocktail supplemented with 500 nM ROSI for 14 days elicited a significant and substantial increase in lipid accumulation, as measured by Nile Red fluorescence normalized to cellular DNA content (Hoechst 33342). After treatment with 10 μM 2,4-DTBP for 14 days, adipogenesis was increased significantly whereas lower doses of 2,4-DTBP or all doses of 2, 6-DTBP did not (Fig. 1A). Induction of adipogenesis by LG100268 was similar to that of 2,4-DTBP (Fig. 1A).

**Figure 1.**
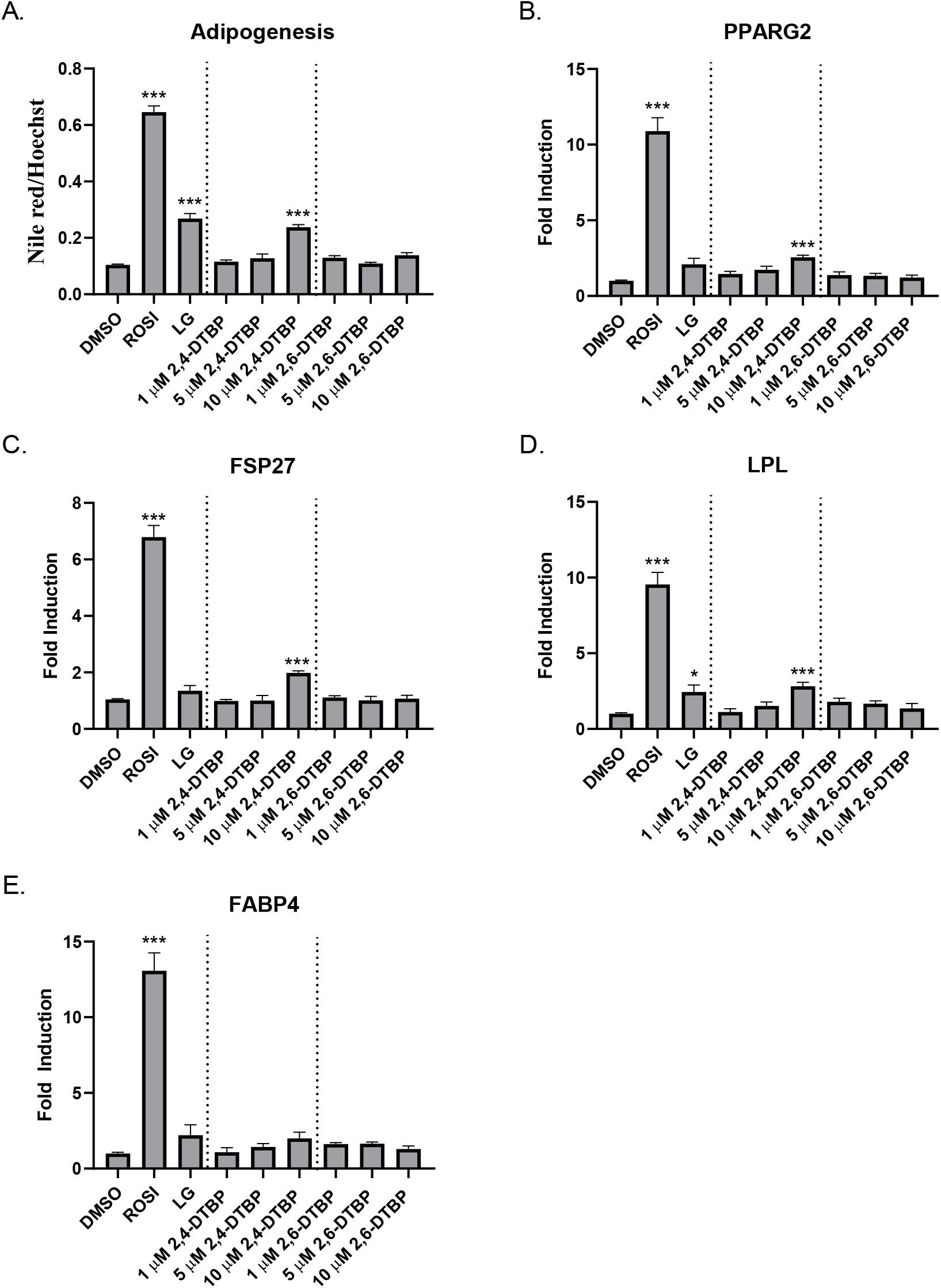
Effect of ROSI, LG100268, 2,4-DTBP, and 2,6-DTBP on lipid accumulation and adipogenic marker gene expression in human MSCs. Cells were treated with 500 nM ROSI, 100 nM LG100268 or different concentrations of 2,4-DTBP or 2,6-DTBP as indicated. Three replicate wells were included for each group in a 24-well plate for lipid accumulation assays or in 12-well plates for adipogenic marker gene expression assay. (A) effects of chemical exposure on adipogenesis in hMSCs. Data are expressed as lipid accumulation normalized to cellular DNA content. (B-E) effects of chemical exposure on expression of adipogenic markers in hMSCs. Data are expressed as fold induction over vehicle control. Error bars represent the standard error of the mean for three replicates. * p ≤ 0.05, ** p ≤ 0.01, *** p ≤ 0.001 compared with cells treated with 0.1% DMSO vehicle control.

We confirmed the adipogenic effects by evaluating the expression of white adipocyte marker genes in response to chemical exposure. Levels of mRNAs encoding FABP4, FSP27, LPL, and PPARγ-2 were significantly increased in the 500 nM ROSI treated cells compared with vehicle (DMSO) (Fig 1B-E). Levels of FSP27, LPL, and PPARγ-2 mRNAs were increased in cells treated with 10 μM 2,4-DTBP compared with 2,6-DTBP and vehicle control, both of which were inactive. Levels of FABP4 were increased, but this increase did not reach statistical significance in 2,4-DTBP-treated cells.

### 2,4-DTBP-increased adipogenesis was blocked by the PPARγ antagonist T0070907 or RXRα antagonist HX531

PPARγ plays a key role in regulating the transcription of genes involved in adipogenesis in MSCs ^31^. Therefore, we explored whether antagonizing PPARγ activation with the specific antagonist, T0070907 affected 2,4-DTBP-mediated increases in lipid accumulation in human MSCs. We found that co-treatment with 10 μM T0070907 ^32^ significantly (*p*<0.05) inhibited lipid accumulation induced by treatment with 100 nM ROSI or by 10 μM 2,4-DTBP (Fig. 2A).

**Figure 2.**
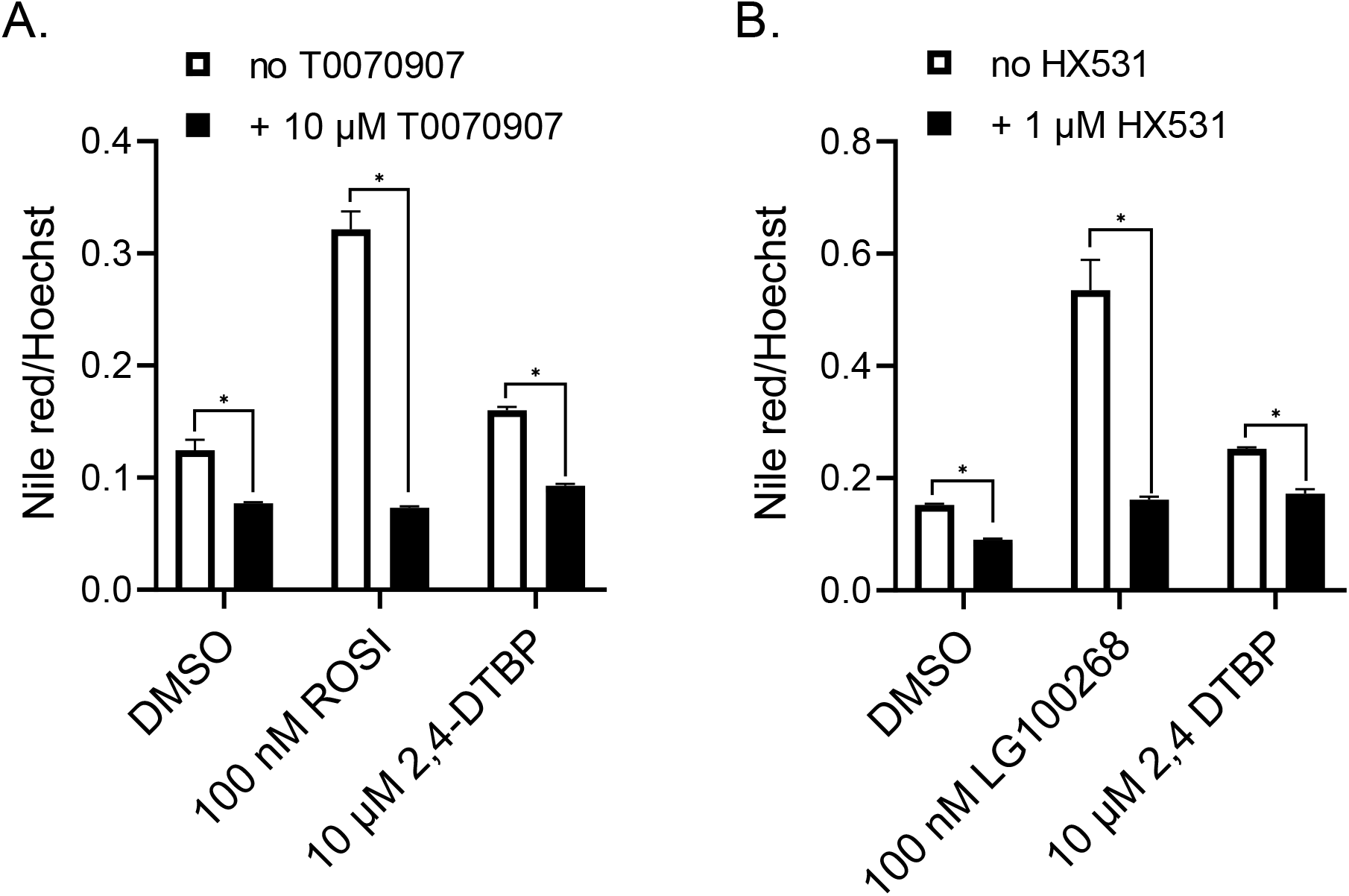
Study of the inhibitory effects of PPARγ antagonist T0070907 and RXRα antagonist HX531 on lipid accumulation induced by 2,4-DTBP in human MSCs. (A) Cells were treated with 100 nM ROSI or 10 μM 2,4-DTBP in the absence or presence of 10 μM T0070907. (B) Cells were treated with 100 nM LG100268 or 10 μM 2,4-DTBP in the absence or presence of 1 μM HX531. Data are expressed as lipid accumulation normalized to cellular DNA content. Error bars represent the standard error of the mean for three replicates. * p ≤ 0.05, ** p ≤ 0.01, *** p ≤ 0.001 compared with cell samples treated with 0.1% DMSO vehicle.10 μM T0070907 or 1 μM HX531.

PPARγ regulates the transcription of adipogenic genes as a heterodimer with RXRs ^31^. To assess the extent of RXR involvement in 2,4-DTBP-induced lipid accumulation, we conducted an RXR antagonist assay by comparing the effects of 2,4-DTBP in the presence or absence of the specific antagonist HX531 ^33^. Co-treatment with 1 μM HX531 significantly (*p*<0.05) inhibited lipid accumulation induced by 100 nM of the specific RXR agonist, LG100268 ^22^ and by 10 μM 2,4-DTBP (Fig. 2B). Taken together, these results implicate the PPARγ-RXR heterodimer in 2,4-DTBP-induced adipogenesis, but not which partner is responsible.

### 2,4-DTBP showed agonistic activity toward GAL4-PPARγ and GAL4-RXRα

We next assessed whether 2,4-DTBP could activate human PPARγ or RXRα using transient transfection in COS-7 cells with GAL4-receptor chimeras. ROSI enhanced the activity of a GAL4-PPARγ in a dose-dependent manner with a lowest effective concentration (LOEC) value of ∼10 nM (Fig. 3A). Similarly, 2,4-DTBP enhanced GAL4-PPARγ activation in a dose-dependent manner with a LOEC value of ∼10 μM (Fig. 3B). 2,6-DTBP did not activate GAL4-PPARγ between 1.2–40 μM (Fig. 3C).

**Figure 3.**
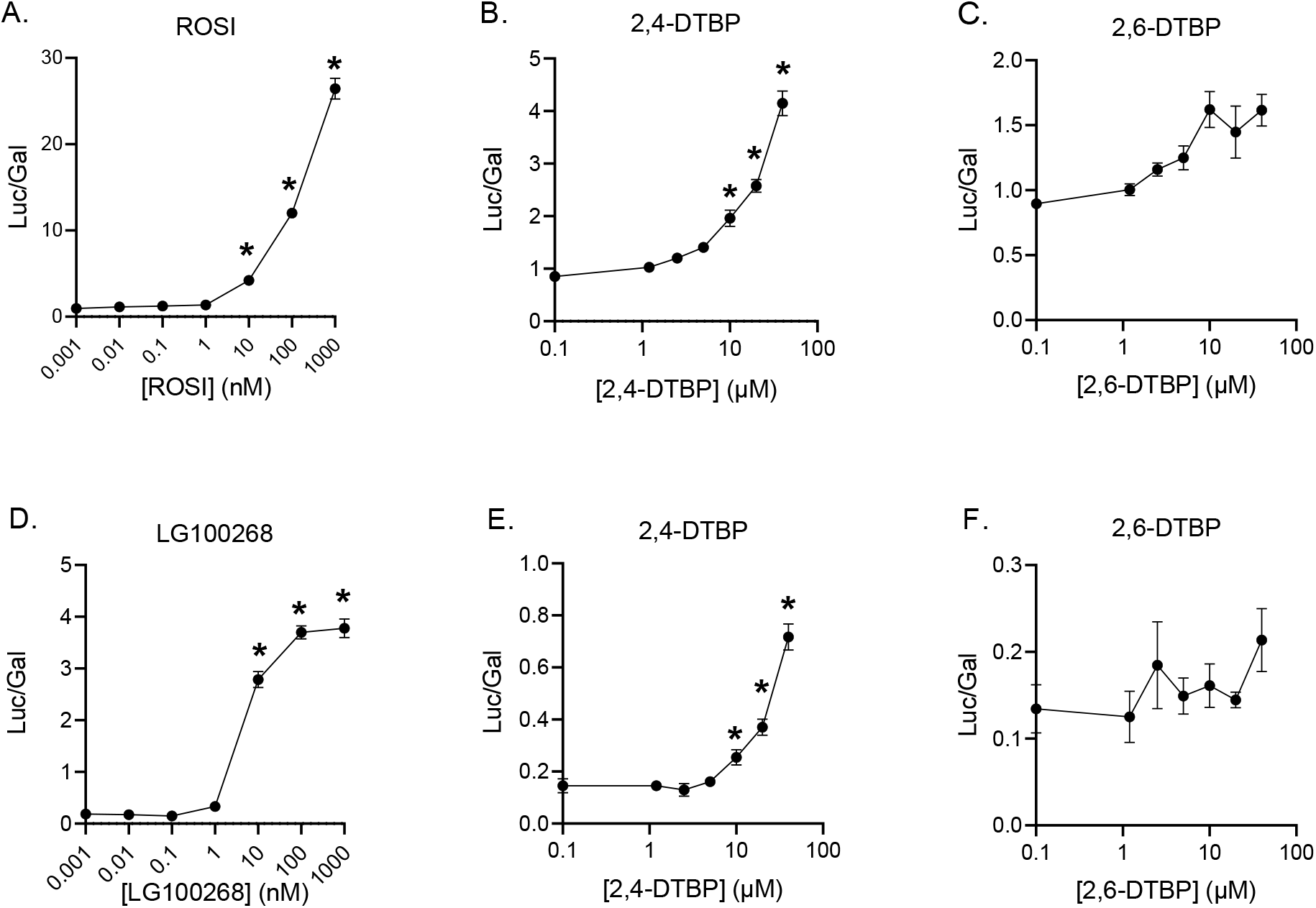
The effect of ROSI, LG100268, 2,4-DTBP, and 2,6-DTBP on GAL4-PPARγ (A-C) and GAL4-RXRα (D-F) regulated luciferase transcriptional activity in COS-7 cells. Plasmid transfected cells were treated with different concentrations of chemicals for 24 h. Three replicated wells were included for each group in a 96-well plate. Data are reported as relative light units (Luc/Gal). Each luciferase read was normalized with the corresponding β-galactosidase read coming from the same transfection well and multiplied by the number of minutes the β-galactosidase plate was incubated (15 min). Error bars represent the standard error of the mean for three replicates. * p ≤ 0.05, compared with cell samples treated with 0.1% DMSO vehicle.

Interestingly, 2,4-DTBP also showed agonistic activity toward human RXRα. Treatment with LG100268 strongly induced activity by GAL4-RXRα in transient transfection assays also with a LOEC of ∼10 nM (Fig. 3D). 2,4-DTBP also enhanced GAL4-RXRα activity with a LOEC value of ∼10 μM (Fig. 3E). 2,6-DTBP was inactive (Fig. 3F). Taken together, these results confirm that 2,4-DTBP appeared to have agonistic activity toward both PPARγ and RXRα.

### RXRα mediated the agonistic activity of 2,4-DTBP toward PPARγ

We next asked whether 2,4-DTBP could interfere with the ability of ROSI to activate GAL4-PPARγ as would be expected by a bona fide PPARγ ligand. We tested the ability of different concentrations of 2,4-DTBP to affect activation of GAL4-PPARγ by 500 nM ROSI and found no significant increase or decrease in activity (Fig. 4A) suggesting that 2,4-DTBP and ROSI did not compete for the same binding site in PPARγ. Since 2,4-DTBP did not further activate GAL4-PPARγ, we inferred that it does not directly interact with PPARγ.

**Figure 4.**
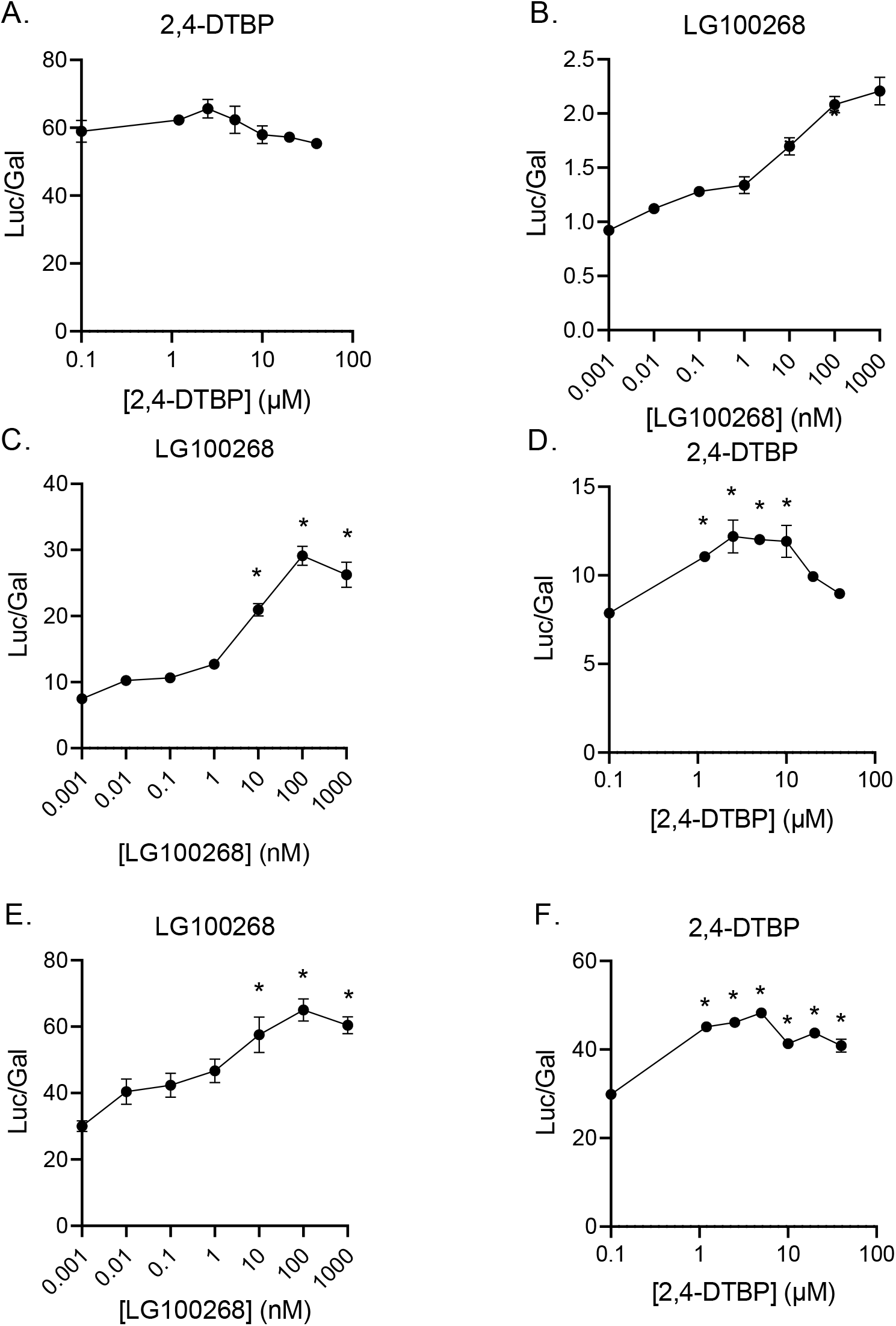
The effect of LG100268 and 2,4-DTBP on GAL4-PPARγ or GAL4-PPARγ/L-RXRα-regulated luciferase activity in COS-7 cells. (A, E, F) ROSI interference assays evaluating the ability chemicals to interfere with ROSI-induced reporter activity on GAL4-PPARγ (A), or GAL4-PPARγ/L-RXRα-induced reporter activity (E, F). (B, C, and D) Agonist assays evaluating the effects of chemicals on ability of GAL4-PPARγ (B), or GAL4-PPARγ/L-RXRα (C, D) to elicit luciferase reporter activity. The concentration of ROSI were 500 nM and 25 nM in the GAL4-PPARγ and GAL4-PPARγ/L-RXRα-regulated luciferase reporter gene assay, respectively. Transfected cells were treated with indicated concentrations of chemicals for 24 h in triplicate in 96-well plates. Data are reported as relative light units (Luc/Gal). Error bars represent the standard error of the mean for three replicates. * p ≤ 0.05, compared with cell samples treated with 0.1% DMSO vehicle. (B, C, and D) or with 500 nM ROSI (A) or 25 nM ROSI (E, F).

Since PPARγ/RXR is a permissive heterodimer, activation of RXR in the heterodimer could lead to activation of the unliganded PPARγ or further enhance the activity of the agonist-bound PPARγ ^23^. Because COS-7 cells express endogenous RXRα ^34^, it is possible that there is some interaction between endogenous full-length RXRα and the GAL4-PPARγ heterodimer in the transient transfection assays. Since 2,4-DTBP had agonistic activity toward RXRα (Fig. 3E), we hypothesized that 2,4-DTBP might activate GAL4-PPARγ via a heterodimer with endogenous RXRs. In support of this hypothesis, we found that LG100268 could also elicit modest activation of GAL4-PPARγ (Fig. 4B).

To further explore the involvement of RXRα in activation of the PPARγ/RXRα heterodimer by 2,4-DTBP, we tested the influence of co-expression of the ligand-binding domain of RXRα (L-RXRα) on the effect of 2,4-DTBP toward the GAL4-PPARγ. We co-transfected the COS-7 cells with two plasmids expressing GAL4-PPARγ and L-RXRα, which lacks the ability to bind DNA and must exert any observed actions via its heterodimeric partner ^23^. Co-transfection with L-RXRα strongly potentiated the ability of LG100268 to activate GAL4-PPARγ (Fig. 4C). Similarly, 2,4-DTBP was now able to induce robust activation of the GAL4-PPARγ-L-RXRα heterodimer at 1 μM (Fig. 4D). Both LG100268 and 2,4-DTBP potentiated reporter gene activity induced by 25 nM ROSI in cells co-transfected with GAL4-PPARγ and L-RXRα although LG100268 potentiation was only significant at 10 nM and higher Fig. 4E, F). We inferred that 2,4-DTBP action on the PPARγ-RXR heterodimer is mediated through RXR since expression of exogenous RXRα ligand binding domain potentiated action of 2,4-DTBP and the RXR-selective agonist LG100268 on GAL4-PPARγ.

### 2,4-DTBP activated GAL4-LXRα and GAL4-TRβ through RXRα in the heterodimers

In addition to PPARγ, RXRα could also heterodimerize with many other nuclear receptors including LXR, TR, RAR, constitutive androstane receptor, farnesoid X receptors, pregnane X receptor, and vitamin D receptor ^35, 36^. Since we found that 2,4-DTBP activation of RXRα in the heterodimers could affect the function of PPARγ, we next asked whether 2,4-DTBP could also modulate the function of other nuclear receptors through activation of RXRα in the heterodimers. To test this hypothesis, we studied the effects of 2,4-DTBP on human GAL4-LXRα and GAL4-TRβ. The results obtained for GAL4-LXRα were similar to those shown for GAL4-PPARγ in Fig. 4. In COS-7 cells transfected with GAL4-LXRα alone, LG100268 (Fig. S1A) and 2,4-DTBP (Fig. S1B) increased reporter gene activity significantly. This increase was strongly potentiated by co-transfection with L-RXRα (Fig. S1E, F). Neither LG100268 (Fig. S1C) nor 2,4-DTBP (Fig. S1D) showed any interference with activation of GAL4-LXRα by its specific agonist GW3965 ^37^. When cells were additionally co-transfected with L-RXRα, activation by GW3965 was potentiated by LG100268 (Fig. S1G) and 2,4-DTBP (Fig. S1H). We inferred that LG100268 and 2,4-DTBP activated GAL4-LXRα heterodimer via its RXRα component, rather than LXRα directly.

We next tested the effects of these chemicals on TRβ and observed that LG100268 (Fig. S2A, C) and 2,4-DTBP (Fig. S2B, D) neither activated nor interfered with activation of GAL4-TRβ by T3. However, co-transfection with L-RXRα allowed LG100268 (Fig. S2E, G) and 2,4-DTBP (Fig. S2F, H) to activate GAL4-TRβ in the absence (Fig. S2E, F) or presence (Fig. S2G, H) of its specific ligand, T3. These results further supported a model in which 2,4-DTBP alters the function of nuclear receptor heterodimers via activation of the RXRα partner.

### Crystallographic analysis of the binding of 2,4-DTBP to RXRα

Prototypical synthetic or natural rexinoids, the subclass of retinoids targeting RXRα such as 9-cis-retinoic acid, contain a carboxylic head group involved in a salt bridge with an arginine residue and a long aliphatic/aromatic chain making numerous van des Waals contacts with essentially non-polar residues distributed all over the ligand-binding pocket (LBP) of RXRα. Obviously, 2,4-DTBP neither structurally nor chemically resembles classical rexinoids. To gain structure-based insight into its mode of action, we solved the crystal structure of the RXRα LBD (hereafter RXR) bound to 2,4-DTBP and a short LxxLL-containing peptide derived from the transcriptional intermediary factor 2 (TIF2), at 2.35 Å resolution (Supplementary Table 1). The structure reveals a RXR homodimer with 2,4-DTBP and the coactivator peptide bound to both protomers (Fig. 5A). Each subunit displayed the canonical active conformation with the C-terminal helix H12 (also termed AF-2 or activation helix) capping the LBP, and the 2,4-DTBP could be precisely placed in its electron density. Interestingly, the density map allowed the unambiguous identification of two alternative orientations of 2,4-DTBP that can be reached through rotation by 180° around the hydroxyl axis (Fig. 5A, inset).

**Figure 5.**
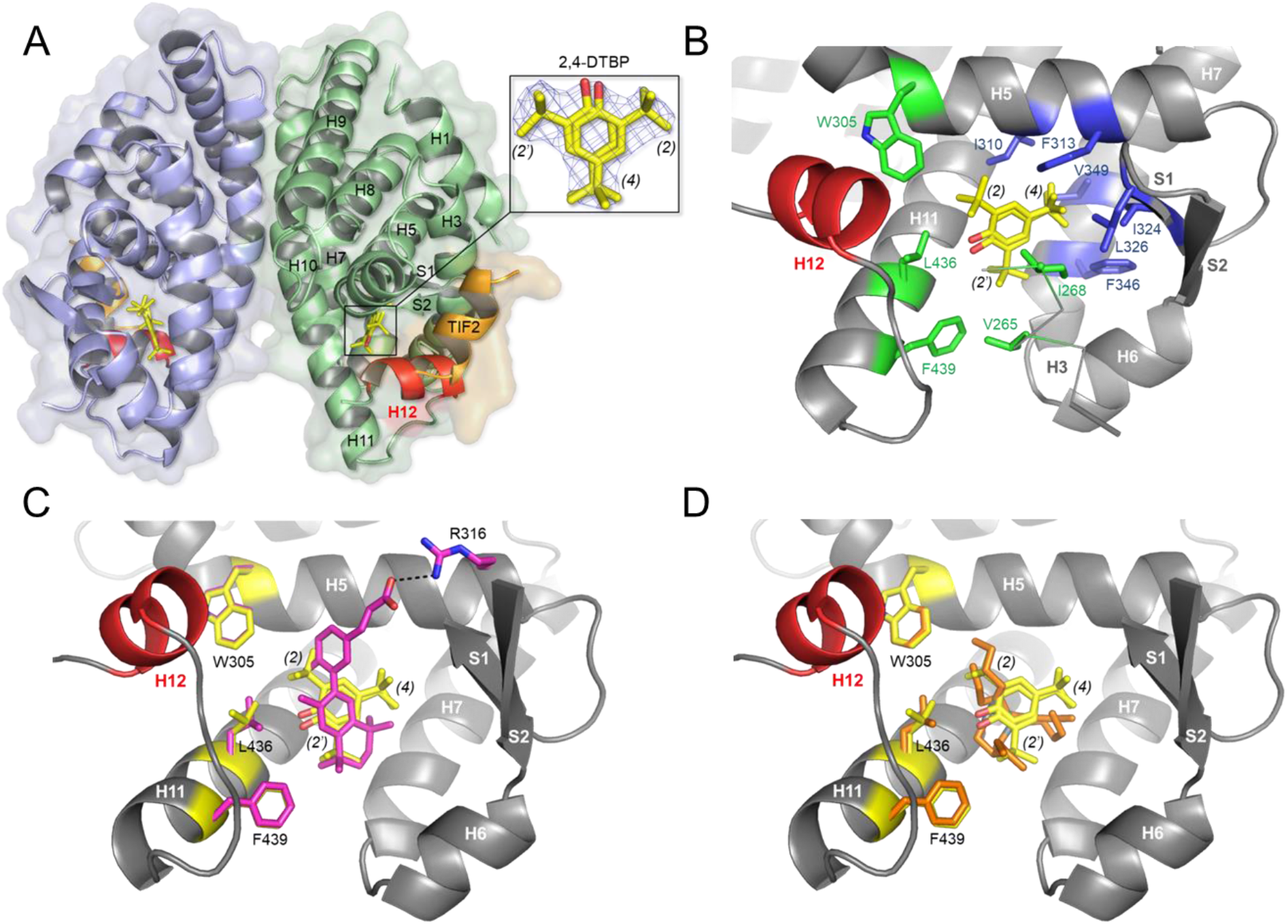
Structural basis for RXR interaction with and activation by 2,4-DTBP. (A) Overall structure of RXR LBD homodimer in complex with 2,4-DTBP (yellow sticks) and TIF2-NR2 peptide (orange). The activation helix H12 is highlighted in red. The inset shows the 2Fo-Fc electron density map of 2,4-DTBP. (B) Close-up view of the ligand-binding pocket of RXR bound to 2,4-DTBP. Key RXR residues in contact with the tert-butyl moieties at positions 2, 2’ and 4 are depicted as green and blue sticks, respectively. (C) Superposition of the 2,4-DTBP- and CD3254-bound RXR structures (PDB code 3E94). As a prototypical rexinoid, CD3254 (magenta sticks) contains an acidic head group and a long aliphatic/aromatic chain bridging arginine 316 (H5) on one side and H11 on the other. (D) Superposition of the 2,4-DTBP- and TBT-bound RXR structures (PDB code 3FUG) showing that the binding sites of the two compounds overlap. TBT is displayed as orange sticks.

2,4-DTBP bound between helices H3 and H11, and in close proximity to C432. While not directly in contact with the AF-2 helix, the tert-butyl group at position 2 (and the alternative 2’ site) plays an important role in stabilizing key residues of helices H3, H5 and H11 to which helix H12 docks in the transcriptionally active conformation (green residues in Fig. 5B). On the other hand, the tert-butyl at position 4 makes extensive van des Waals contacts with hydrophobic residues deeply buried in the LBP (Fig. 5B, residues colored in blue). These interactions are critical for the binding affinity of 2,4-DTBP as well as for its activity, by maintaining the tert-butyl group at position 2 in close proximity to residues involved in receptor activation. The hydroxyl moiety of 2,4-DTBP appears to be involved in a hydrogen bond with the main chain carbonyl group of C432.

Comparison of the present structure with that of RXRα bound to the reference synthetic agonist CD3254 ^38^ shows that 2,4-DTBP occupies only a small fraction of the LBP, but that interactions with residues which are crucial for stabilization of the receptor-active form are preserved in both complexes (Fig. 5C). Interestingly, similar observations were previously made with the potent RXR environmental agonist tributyltin ^39^. Indeed, superposition of the 2,4-DTBP- and TBT-bound RXRα structures revealed that the binding sites of the two compounds largely overlap (Fig. 5D). Overall, it appears that, as for TBT, although 2,4-DTBP interacts with only a subset of binding pocket residues in the H11 region, it is engaged in enough essential contacts to stabilize RXRα in its active conformation. However, the high-affinity binding of TBT is ensured by a covalent bond linking the metal atom of the organotin and cysteine residue 432 in helix H11 (masked by the ligands in Fig. 5D). This covalent bond doesn’t exist for 2,4-DTBP which, as a consequence, binds much less avidly than TBT to RXRα.

### Structure-activity analysis of 2,4-DTBP and congeners on RXRα based on stably transfected HELN RARβ reporter cells

The crystallographic data revealed the structural basis for the agonistic activity of 2,4-DTBP toward RXRα and could allow a better understanding of the binding and activation properties of its congeners. We next investigated the activity of 2,4-DTBP and closely related compounds toward RXRα/RARβ heterodimer by using the stably transfected HELN RARβ reporter cell line in which both RXR and RAR agonist are able to induce luciferase expression. We monitored the agonist potential of 2,4-DTBP and four related compounds (Supplementary Fig. 3A), namely the 2,4,6-tri-tert-butylphenol bearing an additional tert-butyl group at position 6 (2,4,6-TTBP), the 2,6-DTBP containing two tert-butyl groups in ortho positions, and the two corresponding benzene derivatives lacking the hydroxyl moiety, the 1,3-di-ter-butylbenzene (1,3-DTBB) and 1,3,5-tri-tert-butylbenzene (1,3,5-TTBB). We found that 2,4,6-TTBP is, by far, the most potent RXRα agonist, followed by 1,3,5-TTBB and 2,4-DTBP, while 1,3-DTBB and 2,6-DTBP are inactive (Fig. S3A). To assess the specific effect of 2,4-DTBP, 2,4,6-TTBP and 1,3,5-TTBB, HELN RARβ cells were co-incubated with saturating concentrations of the RAR and RXR agonists CD3254 or TTNPB. 2,4-DTBP, 2,4,6-TTBP and 1,3,5-TTBB are able to further activate the TTNPB-saturated RXR-RARβ heterodimer (Fig. S3B). However, 2,4-DTBP, 2,4,6-TTBP and 1,3,5-TTBB appear unable to act in conjunction with CD3254 to enhance the activity of RXR-RARβ (Fig. 3C). These transactivation experiments confirmed the ability of 2,4-DTBP, 2,4,6-TTBP and 1,3,5-TTBB to activate the RXRα-RARβ heterodimer through RXRα. These activity profiles are in full agreement with our structural analysis and confirm that the tert-butyl groups at positions 2, 4 and 6 (2’ site in the 2,4-DTBP complex structure) are important for compound affinity and activity. With three tert-butyl groups, 2,4,6-TTBP displays the best potency and efficacy values followed by 1,3,5-TTBB differing from the latter by the lack of the hydroxyl moiety. The specific role of the tert-butyl at position 4 suggested by the crystal structure was also confirmed by cellular assays since the 2,6-DTBP containing only two tert-butyl groups on both sides of the hydroxyl moiety shows no detectable activity whereas with two tert-butyl groups in ortho and para positions, 2,4-DTBP acts an effective RXRα activator.

## DISCUSSION

2,4-DTBP has been detected in urine samples from the general U.S. population at high concentrations ^18^; however, information about its toxicity is very limited. Since 2,4-DTBP is widely used as an antioxidant in plastics, is known to interact with nuclear hormone receptors ^40^ and has been detected in humans, we asked whether 2,4-DTBP is a potential obesogen and, if so, what is its underlying mechanism of action.

We used an established MSC assay ^19, 20, 22, 32, 41^ to evaluate the potential of 2,4- and 2,6-DTBP for obesogenic activity. We found 2,4-DTBP exposure induced lipid accumulation in human MSCs with a corresponding increase in the induction of marker genes for white adipocyte differentiation (Fig. 1). In contrast, 2.6-DTBP was inactive in these assays suggesting that 2,4-DTBP is a potential obesogen. We further found that 2,4-DTBP can activate the PPARγ-RXRα heterodimer – the so-called “master regulator” of adipogenesis ^31^. Activation of PPARγ is a well-established predictor of adipogenic activity, *in vivo*, and many chemicals have been shown to promote adipogenesis by activating PPARγ (reviewed in ^29, 30^). We used specific PPARγ and RXRα antagonist assays to confirm the requirement for this heterodimer in promoting adipogenesis (Fig. 2). Transient transfection assays suggested that 2,4-DTBP specifically modulated activation of this heterodimer (Fig. 3).

It is well known that the PPARγ/RXRα heterodimer is a permissive heterodimer because it can be activated by ligands for either partner ^36^. Using a series of GAL4-receptor LBD chimeric receptors and co-transfection with the RXRα LBD we showed that 2,4-DTBP specifically activated the RXRα subunit of the PPARγ-RXR, LXRα-RXRα and TRβ-RXR heterodimers (Figs. 3, 4, S1, S2). Previously, studies showed that activating the RXRα in the PPARγ/RXRα heterodimer played an important role in mediating the obesogenic effect of some environmental chemicals. For example, we found that the agrochemical fludioxonil induced adipogenesis in 3T3-L1 preadipocyte cells through PPARγ/RXRα heterodimer by activating RXRα but not PPARγ ^20^. Moreover, we demonstrated that signaling through RXRα subunit of the PPARγ/RXRα heterodimer was critical to commit MSCs to the adipocyte linage ^22^. Combining the results of human MSCs adipogenesis assay and receptor activation assays, we inferred that 2,4-DTBP is a potential obesogen that acts via RXRα in the context of the PPARγ/RXRα heterodimer.

Different from PPARγ/RXRα heterodimer, the TRβ/RXRα was thought to be a non-permissive heterodimer ^23^. However, RXRα can act as a “nonsilent” partner of TR in particular cellular environments (high ratio of coactivator to corepressor), allowing activation of TR signaling by RXR agonists ^42, 43^. The well-studied environmental obesogen, TBT can act as an RXRα agonist to potentiate TR regulated gene expression and resultant morphological programs directed by thyroid hormone signaling *in vitro* and *in vivo* ^44^. Our studies showed that 2,4-DTBP might activate the TRβ/RXRα heterodimer by activating the RXRα partner. Overall, our results demonstrated that the 2,4-DTBP activated the RXRα component of PPARγ/RXRα, LXRα/RXRα, and TRβ/RXR heterodimers.

We next solved the crystal structure of 2,4-DTBP bound to RXRα and identified its specific mode of binding (Fig. 5). The structure explained the RXR binding and activation properties of 2,4-DTBP and allowed us to predict the activity of various congeners. Notably, we predicted that 2,4,6-TTBP would be a more potent activator of RXR, and receptor activation assays confirmed this prediction (Fig. S3).

Because 2,4-DTBP received little attention previously, limited information was available about its toxicity. 2,4-DTBP elicited hepatic toxicity (related to centrilobular hypertrophy of hepatocytes which resulted in heavier livers) and renal toxicity (tubular basophilia and an increased incidence of proteinaceous and granular casts) to rats after oral administration at 300 mg/kg/day for 28 days ^45^. 2,4-DTBP also increased cholesterol and phospholipid in female rats at the dose of 300 mg/kg/day. Susceptibility of newborn rats to 2,4-DTBP was 4-5 times higher than that of young rats. In a 1-generation fertility and repeated dose toxicity study, parental generation rats were administrated 2,4-DTBP orally for 4 weeks before mating and throughout mating, gestation, and lactation. No treatment-related gross lesions and no impairment on the reproductive capability was found in the parental generation animals treated with 2,4-DTBP at dose levels of up to 300 mg/kg/day. However, dietary administration of 2,4-DTBP at 300 mg/kg daily for 13 weeks to the F1 generation rats elicited a toxic effect on the growth rate, which was secondary to reduced diet palatability ^46^. Based on several *in vitro* assays, 2,4-DTBP was shown to be a possible endocrine-disrupting chemical (EDC). 2,4-DTBP bound to the rainbow trout estrogen receptor (ER) ^47^. Using an effect-directed analysis-based approach to identify EDCs in multi-contaminated river sediment, 2,4-DTBP was identified as a new environmental human estrogen receptor ligand by using gas chromatography coupled with mass spectrometry ^13^.

The molecular mechanisms underlying these observations were unclear. Our data showing that 2,4-DTBP can disrupt nuclear receptor function via RXRα and previous studies linking 2,4-DTBP to estrogen receptor signaling ^13^ provide potential clues to identifying mechanisms underlying these toxic effects. There is considerable evidence linking EDCs and obesity and other health problems ^10, 48, 49^. Many EDCs exert their effects by disrupting the functions of nuclear receptors including PPARs, RXRα, TRs, LXRs, and RARs ^10, 50^. PPARγ, RXRs, LXRα, TRβ and RARs play important roles in many physiological and pathological processes such as development, differentiation, and growth ^51^. The ability of 2,4-DTBP and related chemicals to activate RXRα and RXRα heterodimers implicates it as an EDC and indicates that it might exert endocrine-disrupting effects through these and other nuclear receptors. Comprehensive *in vivo* and *in vitro* toxicology studies are warranted to evaluate the potential effects of 2,4-DTBP and related chemicals.

2,4-DTBP concentrations in human urine samples were in the range of 7.3-130 ng/mL (0.03 μM-0.63 μM). 2,4-DTBP promoted adipogenesis in human MSCs at concentrations of ∼10 μM, which is somewhat higher, but not so far from the concentrations observed in limited studies of human urine. Receptor activation studies showed that 2,4-DTBP and 2,4,6-TTBP could activate RXRs in the low micromolar to sub-micromolar range which is an environmentally relevant level according to the limited human biomonitoring data ^18^. Considering the ability of tert butyl phenols to target RXRα in the context of multiple receptor heterodimers, as well as their ability to activate ERS, additional large-scale surveys on the burden to the human body of 2,4-DTBP are warranted.

Taken together, the data presented here demonstrated that 2,4-DTBP belongs to a family of compounds whose potential endocrine-disrupting and obesogenic effects can be strongly modulated by their chemical composition. Structure-activity studies such as the present one could help the rational development of safer antioxidants.

## ASSOCIATED CONTENT

### Supplemental materials

**Figure S1.**
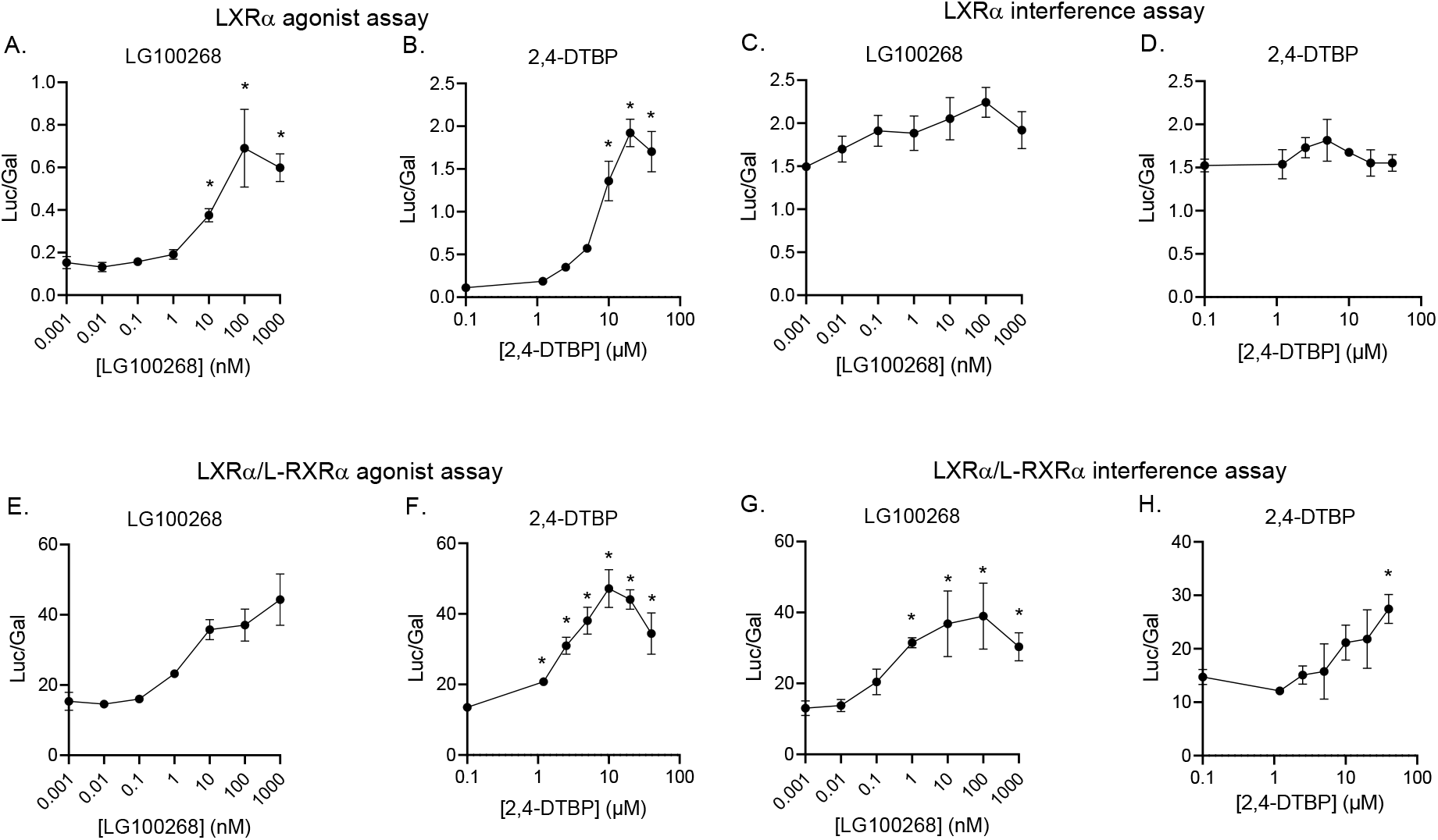
The effects of LG100268 and 2,4-DTBP on GAL4-LXRα- or GAL4-LXRα/L-RXRα-regulated luciferase reporter activity in COS-7 cells. (A, B, E, and F) Agonist assay testing the ability of LG100268 and 2,4-DTBP to induce luciferase assay driven by GAL4-LXRα (A, B) or GAL4-LXRα/L-RXRα (E, F). (C, D, G, H) Interference assay testing the ability of the indicated chemicals to interfere with activation of GAL4-LXRα by 4000 nM (C, D) or GAL4-LXRα/L-RXRα by 400 nM (of the selective LXRα agonist, GW3965. Transfected cells were treated with the indicated concentrations of chemicals in triplicate for 24 hours using 96-well plates. Data are reported as relative light units (Luc/βGal). Error bars represent the standard error of the mean for three replicates. * p ≤ 0.05 compared with cell samples treated with 0.1% DMSO vehicle (A, B, E, and F) or treated with 4000 nM GW3965 (C, D) or 400 nM GW3965 (G, H).

**Figure S2.**
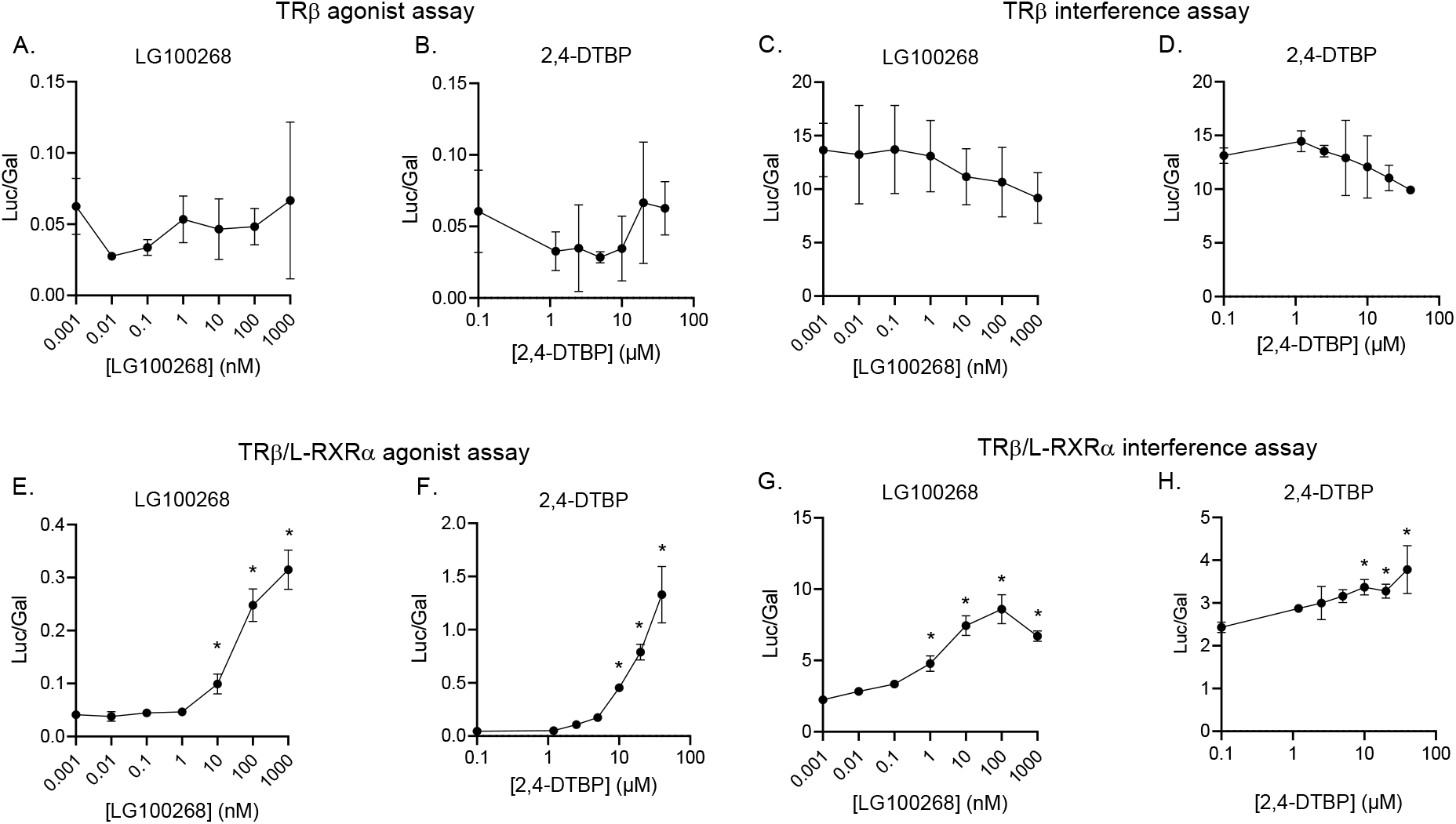
Effects of LG100268 and 2,4-DTBP on GAL4-TRβ or GAL4-TRβ/L-RXRα-regulated luciferase reporter activity in COS-7 cells. Agonist assay testing the ability of LG100268 and 2,4-DTBP to induce luciferase assay driven by GAL4-TRβ (A, B) or GAL4-TRβ/L-RXRα (E, F). (C, D, G, H) Interference assay testing the ability of the indicated chemicals to interfere with activation of GAL4-TRβ by 50 nM of T3 (C, D) or activation by GAL4-TRβ/L-RXRα by 10 nM T3 (G, H). Transfected cells were treated with indicated concentrations of chemicals in triplicate for 24 hours. Data are reported as relative light units (Luc/Gal). Error bars represent the standard error of the mean for three replicates. * p ≤ 0.05, compared with cell samples treated with 0.1% DMSO vehicle (A, B, E, and F). or cell samples treated with 50 nM T3 (C, D) or 10 nM T3 (G, H).

**Figure S3.**
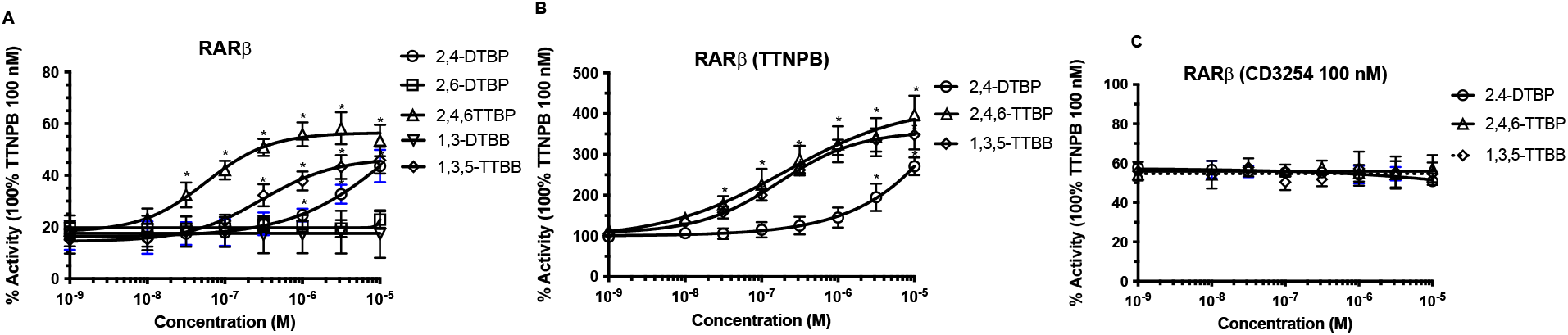
Analysis of tert-butyl phenol and related chemical activation of hRXR. HELN RARβ cells were treated with different concentrations of chemicals for 16 h in absence (A) or presence of the pan-RAR agonist TTNPB (B) or the pan-RXR agonist CD3254 (C). Data are expressed as percentage of the maximal activity obtained with the pan-RAR agonist TTNPB at 100 nM. Error bars represent the standard error of the mean for three experiments. * p ≤ 0.05, compared with cell samples treated with 0.1% DMSO vehicle.

**Table S1.**
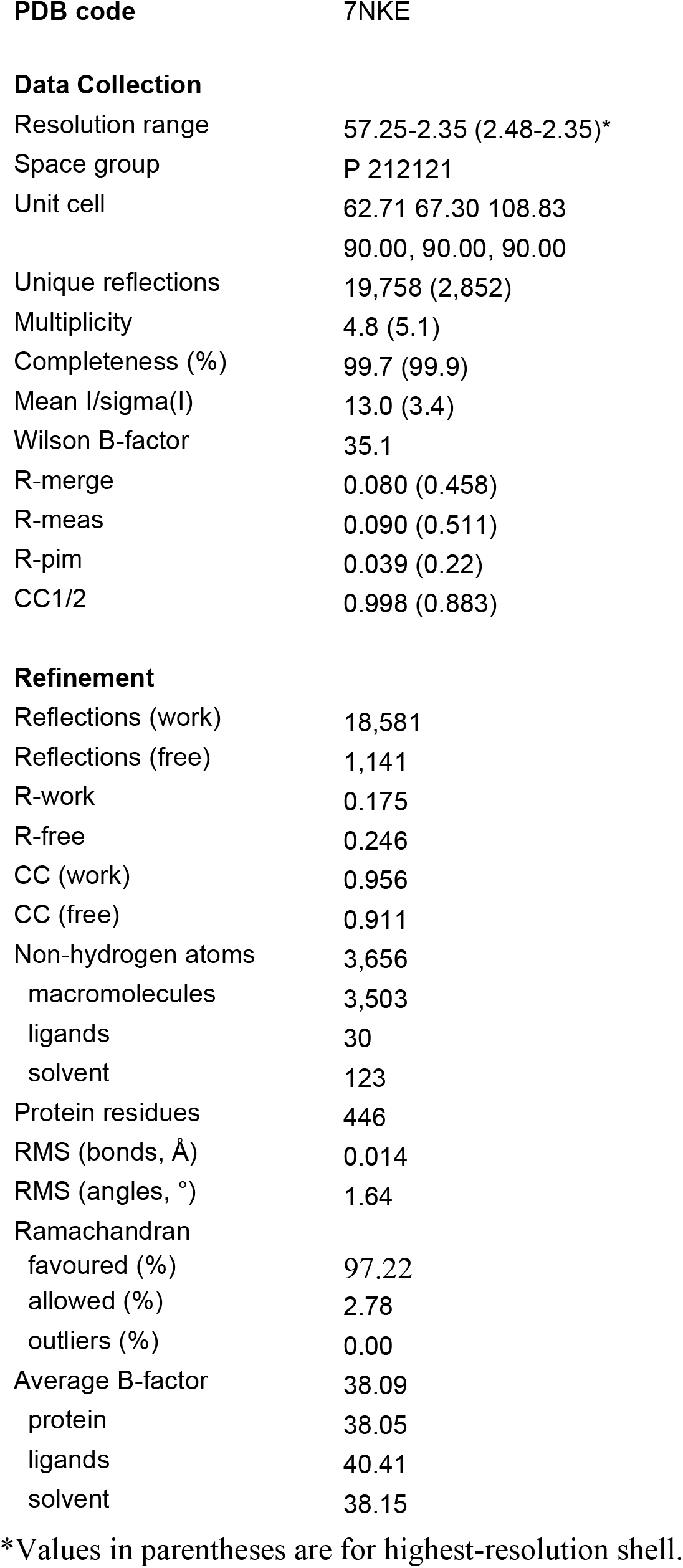
Data collection and refinement statistics

## AUTHOR INFORMATION

### Author Contributions

**Xiao-Min Ren, Richard C. Chang, Yikai Huang, Angela Y. Kuo, Marina Grimaldi, Coralie Carivenc and William Bourguet performed experiments, Xiao-Min Ren, Richard C. Chang William Bourguet, Patrick Balaguer and Bruce Blumberg designed experiments and interpreted data, Xiao-Min Ren, William Bourguet, Patrick Balaguer and Bruce Blumberg wrote the manuscript, William Bourguet, Patrick Balaguer and Bruce Blumberg acquired funding and supervised the experiments. All authors have given approval to the final version of the manuscript**.

## Acknowledgements

We thank Dr. Jane Muncke (Food Packaging Forum Foundation) for suggesting 2,4-DTBP as a compound of interest. We acknowledge experimental assistance from the staff of the European Synchrotron Radiation Facility (Grenoble, France) during crystallographic data collection.

## Abbreviations

DTBP: di-tert-butyl phenol
TTBP: tri-tert-butyl-phenol
DTBB: di-tert-butylbenzene
TTBB: tri-tert-butylbenzene
ROSI: rosiglitazone
RXR: retinoid ‘X’ receptor, aka 9-cis retinoic acid receptor
PPAR: peroxisome proliferator activated receptor
RAR: retinoic acid receptor
BMI: body mass index
SPA: synthetic phenolic antioxidant
BHT: 2,6-di-tert-butyl-4-methylphenol, BHA-3-tert-butyl-4-hydroxyanisosle
GM: geometric mean, BHT
LXR: liver ‘X’ receptor
TR: thyroid hormone receptor
EDC: endocrine disrupting chemical
MSC: multipotent mesenchymal stromal stem cell
FBS: fetal bovine serum
DMSO: Dimethylsulfoxide
QPCR: quantitative real time reverse transcriptase polymerase chain reaction
FABP4: fatty acid binding protein 4
FSP27: fat-specific protein of 27 kDa
PPARγ2: peroxisome proliferator activated receptor gamma 2
LPL: lipoprotein lipase
LBD: ligand-binding domain
LBP: ligand-binding pocket
DBD: DNA-binding domain
EDTA: ethylene diaminetetraacetic acid
DTT: dithiothreitol.

